# Dissection of the corticotroph transcriptome in a mouse model of glucocorticoid-induced suppression of the HPA axis

**DOI:** 10.1101/2020.07.29.227330

**Authors:** N Romanò, PJ Duncan, H McClafferty, O Nolan, Q Ding, NZ Homer, P Le Tissier, BR Walker, MJ Shipston, TJG Chambers

## Abstract

Glucocorticoids (GC) are prescribed for periods >3 months to 1-3% of the UK population; 10-50% of these patients develop hypothalamus-pituitary-adrenal (HPA) axis suppression, which may last over 6 months and is associated with morbidity and mortality. Recovery of higher nodes of the axis is necessary for recovery of adrenal function. We developed a mouse model of Dexamethasone (DEX)-induced HPA axis dysfunction in order to further explore recovery in the pituitary. Adult male C57BL6/J or those crossed with *Pomc*-eGFP mice were randomly assigned to receive DEX (~0.4 mg/kg bodyweight/day) or vehicle via drinking water for 4 weeks following which treatment was withdrawn. Tissues were harvested at 0, 1, and 4 weeks following withdrawal of treatment. Corticotrophs were isolated from *Pomc*-eGFP pituitaries using FACS, and RNA extracted for RNA-seq. DEX treatment suppressed corticosterone production, which remained partially suppressed at least 1 week following DEX withdrawal. In the adrenal, at time 0, *Hsd3b2, Cyp11a1*, and *Mc2r* mRNA levels were significantly reduced, with *Mc2r* and *Cyp11a1* remaining reduced 1 week following DEX withdrawal. The corticotroph transcriptome was modified by DEX treatment with some differences between groups persisting 4 weeks following withdrawal. No genes supressed by DEX exhibited ongoing attenuation 1 and 4 weeks following withdrawal, whilst only 2 genes were upregulated and remained so following withdrawal. A pattern of rebound at 1 and 4 weeks was observed in 14 genes that increased following suppression, and 6 genes that were reduced by DEX and then increased. Chronic GC treatment may induce persistent changes in the pituitary that may influence future response to GC treatment or stress.

## INTRODUCTION

Since the observation of the profound effect of administration of Kendall’s compound E (Cortisone) on the inflammatory condition rheumatoid arthritis (Hench *et al.* 1949), glucocorticoids (GCs) have become a principal treatment for inflammatory conditions across most body systems to the extent that they are now prescribed to 1-3% of the adult population (Fardet *et al.* 2011; Bénard-Laribière *et al.* 2017; Laugesen *et al.* 2017). Despite the evolution of steroid-sparing treatments, prescription rates continue to increase each year (Fardet *et al.* 2011; Laugesen *et al.* 2017).

Exogenous GCs suppress endogenous GC production. When treatment is stopped, homeostasis should re-activate the HPA axis; however, this does not always occur, sometimes with fatal consequences (Fraser *et al.* 1952). Failure of GC production following withdrawal of chronic (>3 month) GC treatment is common (30-50% of people immediately after stopping treatment, 10-20% of people 6 months later (Broersen *et al.* 2015; Joseph *et al.* 2016)). Such cases likely have a significant clinical impact; there is a 2-3x increase in hospital presentations with symptoms and signs of adrenal insufficiency (hypotension, hypovolaemia, cardiovascular collapse, and hypoglycaemia) in the 4 week period following discontinuation of chronic systemic GC treatment (Laugesen *et al.* 2019).

We are unable to accurately identify those patients at most clinical risk; meta-analyses suggest that age, dose, and cumulative dose are likely predictors (Broersen *et al.* 2015; Joseph *et al.* 2016; Laugesen *et al.* 2019) although these only account for some of the variability observed between patients. We also currently have no means of predicting recovery of the HPA axis at the point of starting GC therapy, and thus cannot tailor expensive and involved testing or increased use of steroid-sparing agents to those most at risk of adrenal insufficiency following withdrawal. Certainly, there would be huge clinical benefits to both improve identification of those most at risk and the rate of recovery in those with a suppressed axis.

Chronic GC treatment suppresses all levels of the HPA axis. For example, there are blunted adrenocorticotrophic hormone (ACTH) and cortisol responses to insulin-induced hypoglycaemia (Schlaghecke *et al.* 1992); impaired reactivity of the pituitary to CRH testing (Schürmeyer *et al.* 1985; Schlaghecke *et al.* 1992) and an impaired reactivity of the adrenal to ACTH1-24 (Borresen *et al.* 2017). Importantly, the recovery of ACTH production is essential for recovery of adrenal activity in patients both following treatment for Cushing’s disease and in patients where treatment with supra-physiological exogenous GCs is withdrawn (Graber *et al.* 1965). During recovery there is marked increase in ACTH above physiological levels that precedes recovery of adrenal function.

We established a mouse model to understand the recovery process of the HPA axis following withdrawal of chronic GC treatment. Recovery of the pituitary corticotroph population following withdrawal of chronic GC is essential for recovery of the HPA axis (Graber *et al.* 1965; Hodges & Sadow 1969; Nicholson *et al.* 1984). Having found evidence for disrupted corticotroph function persisting up to one week following withdrawal of GC, and given *in vitro* evidence for long term (120h) transcriptional changes as a result of only a brief (24h) GC exposure (Jubb *et al.* 2017), we hypothesised that chronic GC exposure would program sustained changes in the corticotroph transcriptome. We reasoned that persistent changes to transcriptional regulators or to pathways regulating ACTH synthesis and secretion might explain the delay in corticotroph and thus HPA axis recovery.

## RESULTS

Mice were analysed in 4 groups: those exposed to only drinking water (control; Group A), exposed to dexamethasone (DEX) for 4 weeks (Group B), exposed to DEX for 4 weeks and then switched to drinking water for 1 week (1 week recovery, Group C) or exposed to DEX for 4 weeks and then switched to drinking water for 4 weeks (4 week recovery, Group D)(Figure 1).

**Figure 1.**
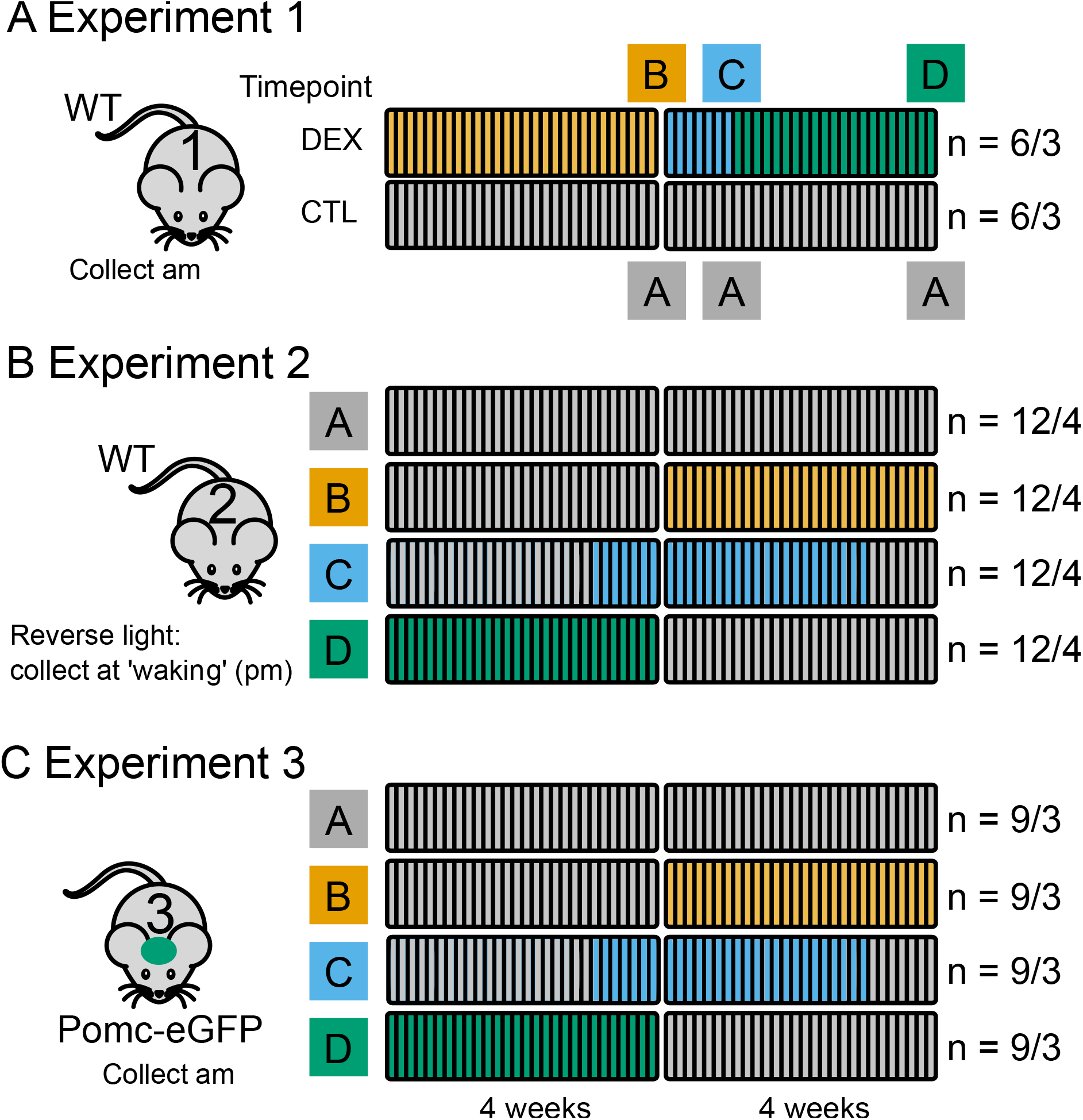
Schematic of experiments. **A. Experiment 1**. C57Bl6/J mice were assigned to receive DEX in drinking water (n = 6 mice in 3 cages) or water alone (n = 6 mice in 3 cages) for 4 weeks. Control mice received only standard drinking water (Group A). Following 4 weeks treatment, 2 animals selected at random from each cage were sacrificed (Group B). Following one further week, two further mice from each cage were collected as at time 0 (Group C). The last animals were collected 4 weeks after DEX withdrawal (Group D). Age matched controls were assigned to group A. **B. Experiment 2.** Here, 16 cages of 3 mice were randomly assigned to 4 treatment groups, A; receiving water in drinking water for 8 weeks, B; drinking water for 4 weeks and then DEX for 4 weeks, C; water for 3 weeks, DEX for 4 weeks then 1 week with drinking water and D; DEX for 4 weeks then drinking water for 4 weeks. All animals were collected at a single time point at the end of the experiment as above. Animals were reverse lit and collected at the point of waking. **C. Experiment 3** was conducted as experiment 2, except that animals were not reverse lit and *Pomc*-eGFP mice were used. 12 cages of 3 animals were assigned randomly to the four groups. At the end of the experiment, tissue was collected as experiment 1 and 2, but anterior pituitary was dissected and dissociated for FACS isolation of corticotrophs. RNA from isolated corticotrophs underwent RNA-seq. Numbers to the right refer to numbers of animals/ in how many cages.

### Body and adrenal weight

Weight gain was attenuated during treatment with DEX. Upon discontinuation of DEX (in group B), weight increased over 28 days to match that of control litter mates. There was a significant interaction between weight, time, and treatment p<0.001 (Figure S1A). Adrenal weight was reduced following treatment and 1 and 4 weeks following withdrawal but this failed to reach the threshold considered significant of p<0.05 (Fig. 2A and 2B and S1B). In experiments 1 and 3, when adrenals were collected at the start of the rest period, this difference remained evident 4 weeks after stopping the DEX. Adrenal/bodyweight ratio was not significantly affected by DEX or withdrawal (Figure 2C and D).

**Figure 2.**
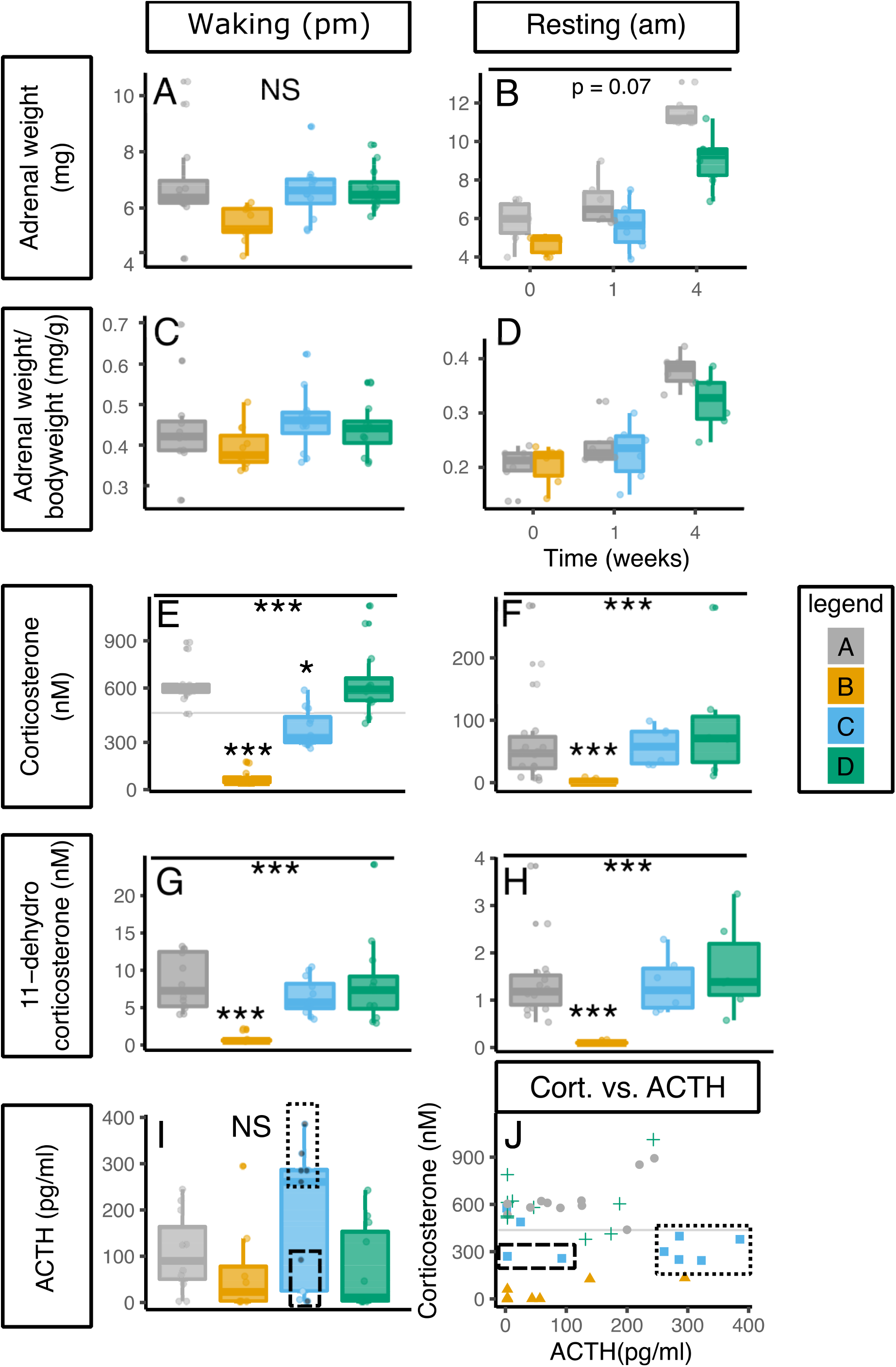
Dexamethasone reduces weight and corticosterone production that persists 1 week following treatment withdrawal. **A and B. Adrenal weight** DEX treatment did not significantly affect adrenal weight. The mean weight of both adrenal glands at collection is presented with box and whisker charts from experiment 2 (A) and experiment 1 (B). **C and D. Adrenal weight/bodyweight D and E.** The adrenal weights as a proportion of bodyweight are presented for experiment 2(C) and experiment 2 (D). **E and F. Corticosterone levels.** DEX treatment reduced corticosterone levels that were still reduced 1 week after stopping DEX treatment at waking (pm) (experiment 2) (E), but which had returned to normal basal levels (am; rest period) (experiment 1) (F). Corticosterone levels had returned to the level of controls 4 weeks after stopping DEX treatment. **G and H 11-dehydrocorticosterone levels.** As E and F but measured 11-dehydrocorticosterone. **I. ACTH levels.** Plasma ACTH was not significantly affected by DEX treatment. One week after stopping DEX, there was greater variation in ACTH levels. Data from experiment 2. The dotted box highlights mice with recovered ACTH production, the dashed box shows mice with unrecovered ACTH production; grey circles in this box are mice with Corticosterone levels below controls, blue circles are mice with corticosterone within the range of controls. **J. Relationship between plasma corticosterone and ACTH.** DEX treatment (yellow triangles) reduced both plasma ACTH and corticosterone compared to controls (grey circles). One week after stopping DEX (blue squares), there was greater variation between animals with some showing high ACTH levels but lower corticosterone (dotted box) and some showing inappropriate ACTH levels for the lower corticosterone levels (dashed box). Four weeks after stopping DEX, ACTH and corticosterone levels had returned to level of controls (green crosses). Data from experiment 2. **Legend.** Grey dots and boxes represent control mice (Group A), yellow those who have had 4 weeks of DEX treatment (Group B), blue those one week after withdrawal of DEX (Group C) and green those 4 weeks after treatment withdrawal (Group D). Data analysed by linear mixed model with group as dependent variable and cage as random factor. Tukey-adjusted post hoc tests compared to control group are indicated above boxes where significant differences were identified. *** p<0.001, ** p<0.01, * p<0.05.

### Steroid levels

Corticosterone levels were significantly reduced by DEX treatment (Fig. 2E and F). When tested at waking – peak corticosterone levels – this finding persisted even 1 week after stopping the DEX (Figure 2 F). 11-Dehydrocorticosterone levels were also reduced by DEX but in both waking and resting phases had recovered 1 week after stopping the DEX (Figure 2G and H). ACTH levels were lower immediately after stopping DEX, though this did not achieve statistical significance. One week after stopping the DEX, two distinct populations of animals were apparent: those with increased ACTH levels (indicating recovery of the higher HPA axis (dotted box)) and those with lower levels than would be expected given the relatively low corticosterone levels (dashed box) (Figure 2F). Two mice had ACTH levels and corticosterone levels consistent with controls (blue circles) (normal corticosterone and low ACTH). This is further exemplified in Figure 2F where the animals in group B (1 week following recovery) form into disparate groups; low ACTH and low corticosterone (dashed box-ongoing HPA axis suppression), high ACTH and low corticosterone (dotted box, recovered pituitary, ongoing adrenal suppression), and normalised ACTH and corticosterone (recovered). DEX levels measured in the plasma from experiment 2 were not detectable in the control group (A) or in group C or D, 1 and 4 weeks after treatment withdrawal, highlighting this was not the reason for lower corticosterone levels in group C. Among those animals exposed to DEX (Group B), plasma DEX levels reached maximally 6nM (Figure S1C).

### Adrenal and hypothalamic gene expression

In experiment 1, DEX treatment had no impact on mRNA detected for *Avp, Crh,* Glucocorticoid receptor (*Nr3c1*), or Mineralocorticoid receptor *(Nr3c2*) in whole hypothalamus (Fig. 3A).

**Figure 3.**
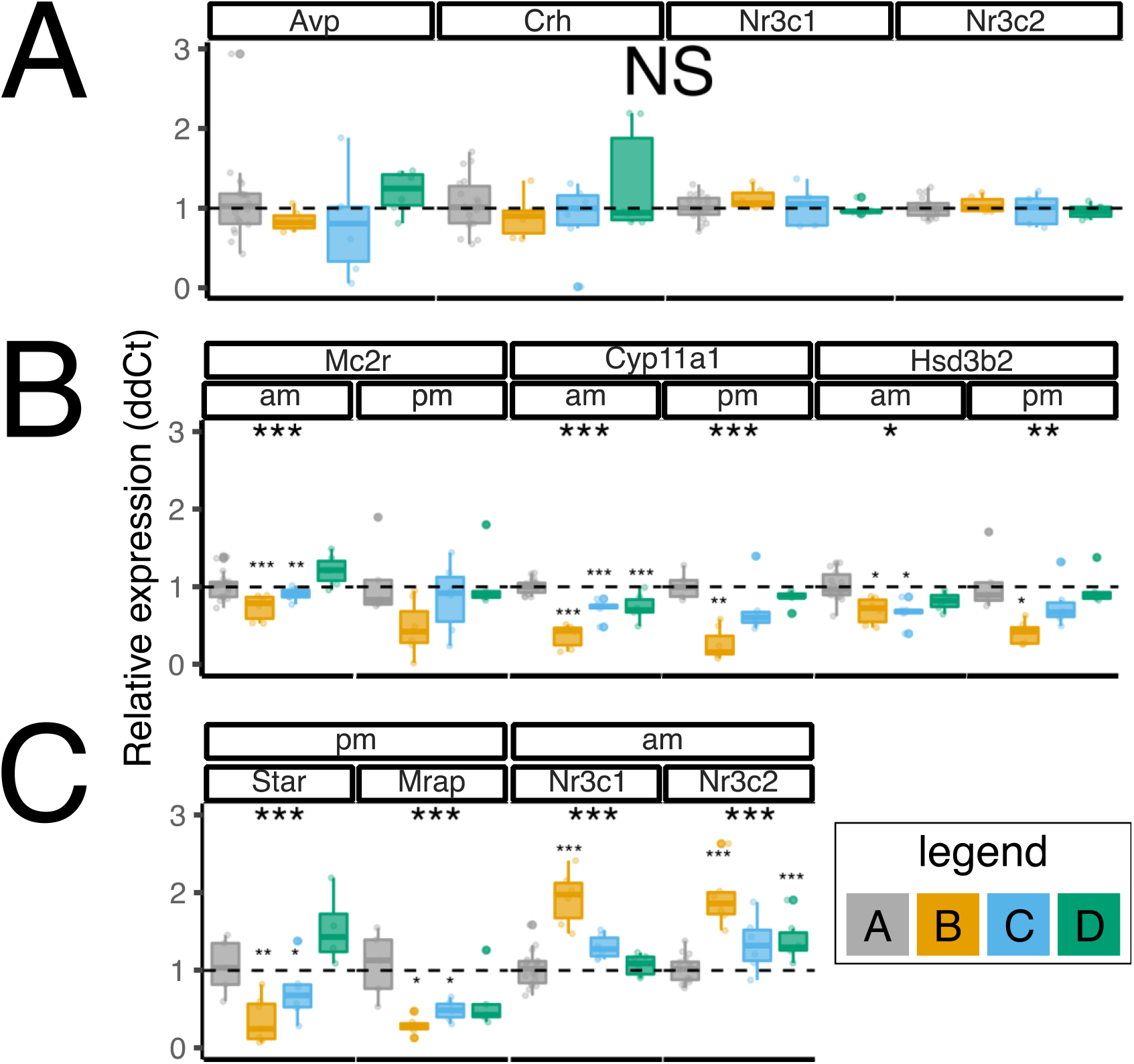
DEX treatment induces significant and persistent changes in adrenal gene expression. **A. Hypothalamus.** DEX treatment did not affect gene expression of *Avp, Crh, Nr3C1*, or *Nr3c2* in whole hypothalamus (data from experiment 1). **B and C. Adrenal.** Dex reduced the adrenal expression of the ACTH receptor (*Mc2r*), steroidogenic enzymes (*Cyp11a1 Hsd3b2* and *Star*) and increased expression of the glucocorticoid receptor (*Nr3c1*) and Mineralocorticoid receptor (*Nr3c2*). Data are shown from experiment 1 (am panels) and experiment 2 (pm panels). For ease of comparison, control samples from each time point in experiment 1 have been combined, ddCt values were made in comparison to time-matched adrenal glands. **Legend.** Grey bars represent control mice (Group A), yellow those who have had 4 weeks of DEX treatment (Group B), blue those one week after withdrawal of DEX (Group C), and green those 4 weeks after treatment withdrawal (Group D). Data analysed by linear mixed model with group as dependent variable and cage as random factor. Tukey-adjusted *post hoc* tests compared to control group are indicated above bars (small asterixis) where significant differences were identified. *** p<0.001, ** p<0.01, * p<0.05.

Gene expression was examined in the adrenal glands from experiments 1 (rest period) and 2 (wake period). DEX treatment significantly reduced expression of mRNA for the ACTH receptor (*Mc2r*), the steroidogenic enzymes *Cyp11a1* (side chain cleavage enzyme) and *Hsd3b2* (3-beta hydroxylase), the first two steps in the steroidogenic pathway, *Star* the gene coding the steroid acute regulatory protein, and the gene coding *Mrap*, which transports the ACTH receptor to the plasma membrane. One week following DEX withdrawal, there was ongoing reduced expression of *Mc2r*, *Mrap*, *Cyp11a1*, and *Star* mRNA. The reduction in mRNA for these genes was greatest in the pm/wake period when, in control animals (group A) corticosterone levels were the highest. Thus, changes to mRNA were present in the adrenal which persist at least one week after withdrawing DEX treatment, consistent with the lower corticosterone levels measured at waking in this group. mRNA levels of the glucocorticoid and mineralocorticoid receptors were significantly increased as a result of DEX treatment, remaining significantly elevated even four weeks after DEX withdrawal in the case of the mineralocorticoid receptor (*Nr3c2*).

### Pituitary gene expression

In humans, following withdrawal of supraphysiological GC, the recovery of adrenal responsiveness to ACTH follows recovery of pituitary ACTH production (presumably a result of the trophic effect of ACTH on the adrenal). In fact, there tends to be an overshoot in ACTH production (as would be seen in primary adrenal failure) prior to recovery of the adrenal (Graber *et al.* 1965). Sustained changes to transcription of the GC responsive gene *Fkbp5* persisting 120h beyond GC withdrawal have been found *in vitro* (Jubb *et al.* 2017). We wished to explore whether transcriptional dynamics were persistently affected following DEX withdrawal and might associate with HPA axis dysfunction. We took advantage of the *Pomc*-eGFP transgenic mouse line to isolate *Pomc* expressing cells from the anterior pituitary in order to assess the corticotroph transcriptome during DEX treatment and following treatment withdrawal (experiment 3). Pilot studies showed ongoing suppression of *Pomc* mRNA in isolated corticotrophs 1 week following DEX withdrawal. We were interested in identifying if other genes showed sustained changes following withdrawal of DEX.

The eGFP expression as measured at the time of FACS exhibited in general a bimodal distribution, suggesting the existence of two sub-populations of *Pomc* expressing cells sorted from the anterior pituitary (Fig. 4A). DEX induced a shift of cells to the lower eGFP-expressing population, which was most apparent four weeks after DEX withdrawal.

**Figure 4.**
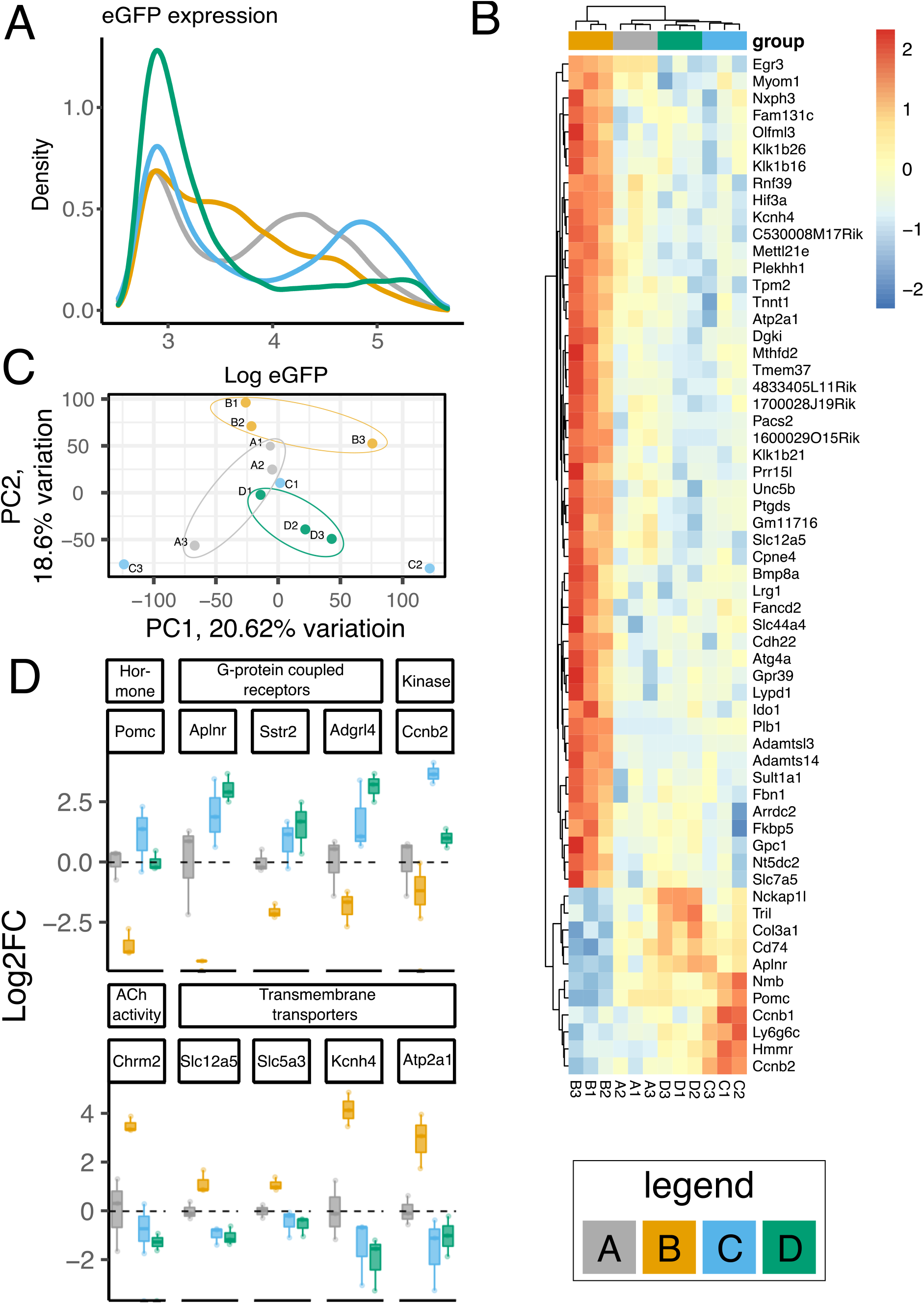
DEX treatment significantly affects the corticotroph transcriptome with changes evident 4 weeks following treatment withdrawal. **A. eGFP expression.** The eGFP expression of the isolated cells is shown. In groups A (control; grey line) and C (1 week post DEX withdrawal; blue line) bimodal populations of cells are apparent. The mean fluorescence of three sorts is shown each comprising the dissociated pituitaries of 2-3 mice. **B. Heatmap showing transcriptomic analysis of isolated cells.** The 60 genes with the most significant difference between groups are shown. Relative expression across each row is shown as per the colour code to the right of the panel. Columns indicate each sample comprising cells isolated from 3 pituitaries and are clustered according to gene expression pattern. Samples are clustered by Euclidean distance. **C. Selected genes from RNA-seq from classes exhibiting persisting changes following withdrawal of DEX.** Log2 fold change data are shown for genes that were significantly affected by treatment and recovery along with the molecular function class to which they belong (see table 1 and 2). Data are the same as those presented in 4B. **Legend.** Grey lines/bars represent control mice (Group A); yellow those who have had 4 weeks of DEX treatment (Group B); blue those one week after withdrawal of DEX (Group C); and green those 4 weeks after treatment withdrawal (Group D).

Treatment with DEX had no impact on the abundance of isolated cells (Fig.S2C), although it did induce a small but not statistically significant reduction in the proportion of GFP-positive cells. RNA-seq confirmed a reduction in *Pomc* mRNA in response to DEX (Fig 4C). Interestingly, 1 week following treatment withdrawal (Group C; green), there was variability in *Pomc* mRNA with increased *Pomc* mRNA in 2 pools, but the third pool had returned to control levels (Figure 4B and 4D).

Of a total of 23,247 genes (with >0.1 read/million in>3 samples), 151 were differentially expressed between the four groups (using cut off false discovery rate <0.05). Primary component analysis showed that control, DEX treatment, and 4-week recovery animals roughly clustered together (Fig 4C) but Group C were dispersed by both PC1 and PC2. The 60 genes exhibiting the most significant difference between groups are shown in Figure 4B. Interestingly all four groups had a distinct transcriptomic signature, particularly remarkable for group D despite this group showing recovery of the HPA axis by measurement of steroidogenesis. DEX treatment resulted in 63 significantly up- and 12 significantly down-regulated genes (log2 fold change >2).

We were next interested in genes that showed persistent changes. We used our dataset of 151 significantly changed genes (FDR <0.05) to find genes that showed persistent changes with log2 fold change of 1 vs. controls (group A). Only small groups of genes showed a persistent change in expression as a result of DEX exposure. 1 gene (*Igha*) was suppressed by DEX and was also suppressed 1 week after treatment withdrawal and 8 genes were both upregulated by DEX and remained elevated one week after withdrawal (*Plb1 Krt19 Fbn1, Ido1, Penk, Klk1b24, Oaf*, and *Col8a2)* two of these remained increased four weeks DEX post withdrawal (*Plb1* and *Krt19*) (Table 1). The majority of differentially expressed genes were increased by DEX and then returned to normal (73 genes) at 1 and 4 weeks.

**Table 1.** 151 differentially expressed genes and the time points at which they were up- or down-regulated. Genes were clustered into 27 subsets based upon their expression relative to controls in each treatment group. Vs. Group B and Group C in rows and Group D in columns. Very few genes demonstrate persistent change in expression in the same direction over all three time points.

| Table 1 |  |  |  |  |
| --- | --- | --- | --- | --- |
| B/D | up | ns | down | D/C |
| up | **group 1**<br>Plb1, Krt19 | **group 7**<br>Adamts14, Fancd2, Klk1b4 | **group 3** | up |
|  | Fbn1, Ido1, Penk, Klk1b24, Oaf, Col8a2 | Ptgs, Klk1b21, Tmem37, Dgki, Klk1b26, 4833405L11Rik, Cdh22, Sult1a1, Pacs2, Gpc1, Olfm13, Nt5dc2, Prr15l, Atg4a, Gpr39, Lypd1, Arrdc2, Tnnt1, Klk1b16, Lrg1, C530008M17Rik, Slc44a4, Cpne4, Slc7a5, Fkbp5, Mthfd2, Unc5b, Gpr1, 1700028J19Rik, Fam131c, Pik3ip1, Ttyh1, Ccnb1, Usp54, Plekhh1, Ciart, E230001N04Rik, Ankmy1, Glb1l2, Nxph3, Aatk, Cobl, Smoc2, Ankdd1b, Mindy4b-ps, Frmd4a, Sez6l, Atp7b, Gm38619, Lrrc75b, Sipa1l1, Angptl7, Gm27252, Sec14l4, Platr25, Unc5a, Slc5a3, Cab39l, Mapt, Ptpv, Wdr93, Rabep2, Fstl3, Prss57, Slc27a3, Susd4, 2410002F23Rik, Slc24a4, Mrps6, Frmpd1 | Adamtsl3, Mettl21e, Bmp8a, Slc12a5, Myom1, Gm11716, Klk1b5, Myh11, Dgkk, Asic2, Pkd1l2, Gm38304 | ns |
|  |  | Atp2a1, Map3k6, Cntfr, Olog | Kcnh4, Hif3a, Tpm2, Rnf39, 1600029O15Rik, Chrm2 | down |
| ns | **group 5**<br>Hmnr, Col4a1, Anln, Col4a2 | **group 8**<br>Ttyh1, Ccnb1 | **group 6** | up |
|  | Tril, Cpxm2, H6pd, Prelp, F2r, Fbln5, Plat, Nid2, Mlc1 | Tmem145, Srpkl, Zbtb16, Cxcl12, Rptor |  | ns |
|  |  | Inhbb | Egr3 | down |
| down | **group 4**<br>Ly6g6c, Aplnr, Pbk, Top2a, Adgrl4, Il16, Plbd1, Kdr, Pdgrfb, Sstr2, Rrm2, H2-Q6, Itgal, Pecam1 | **group 9**<br>Nmb, Ccnb2, Pomc | **group 2**<br>Igha | up |
|  | Col3a1, Cd74, Nckap1l, H2-Aa, H2-DMb1, H2-Eb1, H2-Ab1, Mfap4 | Psmb8, Rtn4r1l |  | ns |

Another pattern of ‘rebound’ expression was observed, group 4 and group 3 in Table 1. These groups may represent genes that were directly or indirectly stimulated or repressed by GC. 17 genes were upregulated by DEX and then suppressed at 1 or 4 weeks, and 6 genes were upregulated by DEX and then suppressed.

Ontological analysis using Protein Analysis Through Evolutionary Relationships, http://pantherdb.org (Mi *et al.* 2019) identified transporter activity (e.g. GO:0005215; GO:0015075) and kinase activity (GO:0016301) as the molecular processes most affected by DEX (Table 2). Dividing these genes into those up- or down-regulated by DEX further highlighted some classes of molecular function of importance: genes with peptidase/ hydrolase were enriched in those down-regulated by DEX, in particular a number of Kalikrein genes, along with the transporters identified above (Supplemental table 1). Of those suppressed by DEX, antigen binding, extra cellular matrix components, and G-protein coupled receptors were significant (Supplemental table 2).

**Table 2.** Over-representation test for classes of genes with significant differential expression in experiment 3. Genes with differential expression were assessed for over-enrichment using PANTHER version 15.0, GO-slim molecular function. Binomial test with Bonferroni correction was applied to compare the list of differentially expressed genes with that expected.

| PANTHER GO-Slim Molecular Function | Mus musculus - REFLIST | Number of genes (154) | Number of expected genes | Over or under representation | A vs B t (fold Enrichment) | (P-value) |  |
| --- | --- | --- | --- | --- | --- | --- | --- |
| transporter activity (GO:0005215) | 782 | 10 | 5.41 | + | 1.85 | 0.05 | Atp7b Slc7a5 Ttyh1 Slc24a4 Atp2a1 Slc12a5 Slc27a3 Pkd1l2 Slc5a3 Kcnh4 |
| ion transmembrane transporter activity (GO:0015075) | 581 | 9 | 4.02 | + | 2.24 | 0.02 | Atp7b Slc7a5 Ttyh1 Slc24a4 Atp2a1 Slc12a5 Pkd1l2 Slc5a3 Kcnh4 |
| inorganic molecular entity transmembrane transporter activity (GO:0015318) | 550 | 9 | 3.8 | + | 2.37 | 0.01 | Atp7b Slc7a5 Ttyh1 Slc24a4 Atp2a1 Slc12a5 Pkd1l2 Slc5a3 Kcnh4 |
| kinase activity (GO:0016301) | 572 | 8 | 3.96 | + | 2.02 | 0.05 | Srpkl Ccnb2 Dgki Dhkk Ccnb1 Pdgrfb Cab39l Map3k6 |
| phosphotransferase activity, alcohol group as acceptor (GO:0016773) | 534 | 8 | 3.69 | + | 2.17 | 0.03 | Srpkl Ccnb2 Dgki Dgkk Ccnb1 Pdgrfb Cab39l Map3k6 |
| metal ion transmembrane transporter activity (GO:0046873) | 310 | 7 | 2.14 | + | 3.26 | 0.01 | Atp7b Slc24a4 Atp2a1 Slc12a5 Pkd1l2 Slc12a5 Kcnh4 |
| cation transmembrane transporter activity (GO:0008324) | 424 | 7 | 2.93 | + | 2.39 | 0.03 | Atp7b Slc24a4 Atp2a1 Slc12a5 Pkd1l2 Slc5a3 Kcnh4 |
| inorganic cation transmembrane transporter activity (GO:0022890) | 395 | 7 | 2.73 | + | 2.56 | 0.02 | Atp7b Slc24a4 Atp2a1 Slc12a5 Pkd1l2 Slc5a3 Kcnh4 |
| active transmembrane transporter activity (GO:0022804) | 218 | 5 | 1.51 | + | 3.32 | 0.02 | Atp7b Slc24a4 Atp2a1 Slc12a5 Slc5a3 |
| receptor regulator activity (GO:0030545) | 255 | 5 | 1.76 | + | 2.83 | 0.03 | Lypd1 Il16 Inhbb Bmp8a Dkk3 |
| monovalent inorganic cation transmembrane transporter activity (GO:0015077) | 261 | 5 | 1.81 | + | 2.77 | 0.04 | Slc24a4 Atp2a1 Slc12a5 Slc5a3 Kcnh4 |
| active ion transmembrane transporter activity (GO:0022853) | 156 | 5 | 1.08 | + | 4.63 | 0.00 | Atp7b Slc24a4 Atp2a1 Slc12a5 Slc5a3 |
| organic cyclic compound binding (GO:0097159) | 1680 | 4 | 11.62 | - | 0.34 | 0.01 | Chrm Ciart Mrps6 Gata2 |
| kinase binding (GO:0019900) | 158 | 4 | 1.09 | + | 3.66 | 0.02 | Ccnb2 Nell2 Myom1 Ccnb1 |
| heterocyclic compound binding (GO:1901363) | 1638 | 4 | 11.33 | - | 0.35 | 0.01 | Chrm2 Ciart Mrps6 Gata2 |
| calcium ion transmembrane transporter activity (GO:0015085) | 84 | 3 | 0.58 | + | 5.16 | 0.02 | Slc24a4 Atp2a1 Pkd1l2 |
| cytokine activity (GO:0005125) | 116 | 3 | 0.8 | + | 3.74 | 0.05 | Il16 Inhbb Bmp8a |
| nucleic acid binding (GO:0003676) | 1293 | 3 | 8.94 | - | 0.34 | 0.02 | Ciart Mrps6 Gata2 |
| carboxypeptidase activity (GO:0004180) | 30 | 2 | 0.21 | + | 9.64 | 0.02 | Cpxm2 Mindy4b-ps |
| postsynaptic neurotransmitter receptor activity (GO:0098960) | 49 | 2 | 0.34 | + | 5.9 | 0.05 | Chrm2 Lypd1 |
| growth factor binding (GO:0019838) | 38 | 2 | 0.26 | + | 7.61 | 0.03 | Gpc1 |
| glycosaminoglycan binding (GO:0005539) | 42 | 2 | 0.29 | + | 6.88 | 0.03 | Nell2 Rtn4rl1 |
| acetylcholine receptor activity (GO:0015464) | 28 | 2 | 0.19 | + | 10.33 | 0.02 | Chrm2 Lypd1 |
| cyclin-dependent protein serine/threonine kinase regulator activity (GO:0016538) | 35 | 2 | 0.24 | + | 8.26 | 0.02 | Ccnb1 Ccnb2 |
| acetylcholine binding (GO:0042166) | 28 | 2 | 0.19 | + | 10.33 | 0.02 | Chrm2 Lypd1 |
| neuropeptide binding (GO:0042923) | 21 | 2 | 0.15 | + | 13.77 | 0.01 | Sstr1 Gpr1 |
| heparin binding (GO:0008201) | 27 | 2 | 0.19 | + | 10.71 | 0.02 | Nell2 Rtn4rl1 |
| fibroblast growth factor binding (GO:0017134) | 6 | 1 | 0.04 | + | 24.1 | 0.04 | Gpc1 |
| lipopolysaccharide binding (GO:0001530) | 7 | 1 | 0.05 | + | 20.65 | 0.05 | Tril |

**Table 3.**
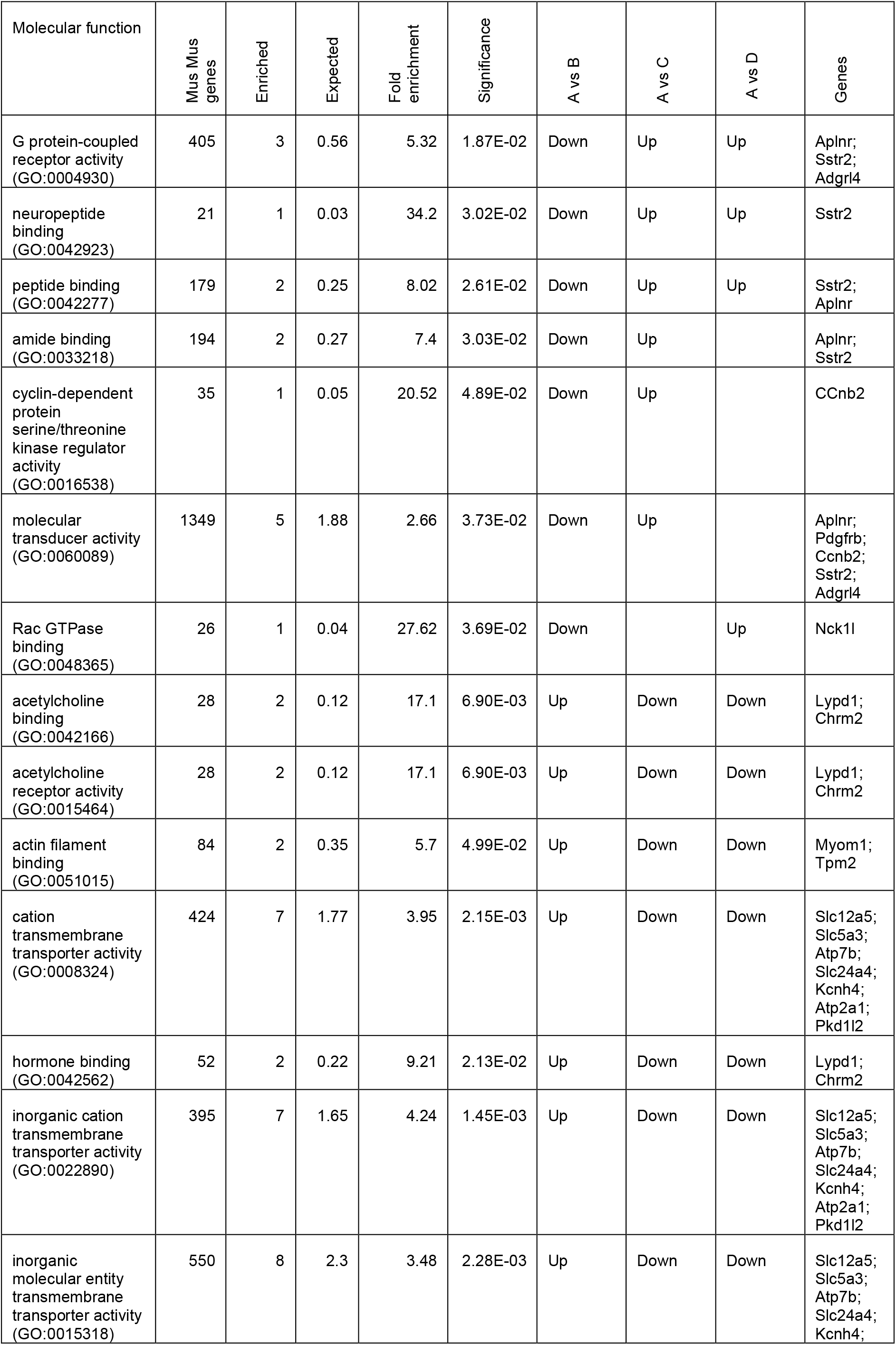
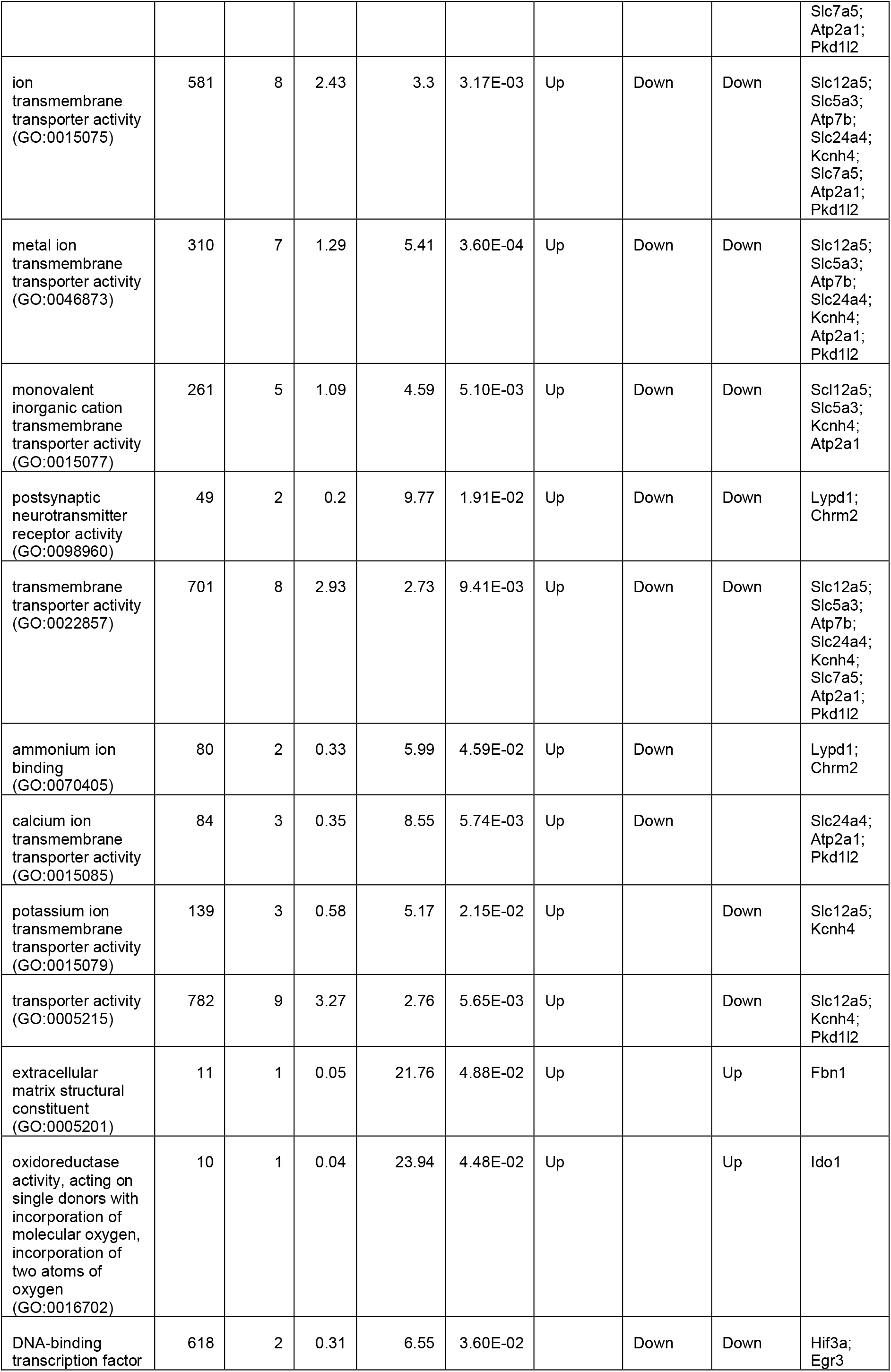
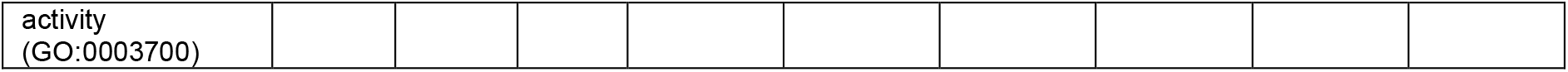
Over-representation test for classes of genes with changes across three groups with direction of change in expression. Genes with differential expression were assessed for over-enrichment using PANTHER version 15.0, GO-slim molecular function. Binomial test with Bonferroni correction was applied to compare the list of differentially expressed genes with that expected. Data are shown from classes identified in more than one comparison.

| Molecular function | Mus Mus genes | Enriched | Expected | Fold enrichment | Significance | A vs B | A vs C | A vs D | Genes |
| --- | --- | --- | --- | --- | --- | --- | --- | --- | --- |
| G protein-coupled receptor activity (GO:0004930) | 405 | 3 | 0.56 | 5.32 | 1.87E-02 | Down | Up | Up | Aplnr;<br>Sstr2;<br>Adgrl4 |
| neuropeptide binding (GO:0042923) | 21 | 1 | 0.03 | 34.2 | 3.02E-02 | Down | Up | Up | Sstr2 |
| peptide binding (GO:0042277) | 179 | 2 | 0.25 | 8.02 | 2.61E-02 | Down | Up | Up | Sstr2;<br>Aplnr |
| amide binding (GO:0033218) | 194 | 2 | 0.27 | 7.4 | 3.03E-02 | Down | Up |  | Aplnr;<br>Sstr2 |
| cyclin-dependent protein serine/threonine kinase regulator activity (GO:0016538) | 35 | 1 | 0.05 | 20.52 | 4.89E-02 | Down | Up |  | CCnb2 |
| molecular transducer activity (GO:0060089) | 1349 | 5 | 1.88 | 2.66 | 3.73E-02 | Down | Up |  | Aplnr;<br>Pdgfrb;<br>Ccnb2;<br>Sstr2;<br>Adgrl4 |
| Rac GTPase binding (GO:0048365) | 26 | 1 | 0.04 | 27.62 | 3.69E-02 | Down |  | Up | Nck1l |
| acetylcholine binding (GO:0042166) | 28 | 2 | 0.12 | 17.1 | 6.90E-03 | Up | Down | Down | Lypd1;<br>Chrm2 |
| acetylcholine receptor activity (GO:0015464) | 28 | 2 | 0.12 | 17.1 | 6.90E-03 | Up | Down | Down | Lypd1;<br>Chrm2 |
| actin filament binding (GO:0051015) | 84 | 2 | 0.35 | 5.7 | 4.99E-02 | Up | Down | Down | Myom1;<br>Tpm2 |
| cation transmembrane transporter activity (GO:0008324) | 424 | 7 | 1.77 | 3.95 | 2.15E-03 | Up | Down | Down | Slc12a5;<br>Slc5a3;<br>Atp7b;<br>Slc24a4;<br>Kcnh4;<br>Atp2a1;<br>Pkd1l2 |
| hormone binding (GO:0042562) | 52 | 2 | 0.22 | 9.21 | 2.13E-02 | Up | Down | Down | Lypd1;<br>Chrm2 |
| inorganic cation transmembrane transporter activity (GO:0022890) | 395 | 7 | 1.65 | 4.24 | 1.45E-03 | Up | Down | Down | Slc12a5;<br>Slc5a3;<br>Atp7b;<br>Slc24a4;<br>Kcnh4;<br>Atp2a1;<br>Pkd1l2 |
| inorganic molecular entity transmembrane transporter activity (GO:0015318) | 550 | 8 | 2.3 | 3.48 | 2.28E-03 | Up | Down | Down | Slc12a5;<br>Slc5a3;<br>Atp7b;<br>Slc24a4;<br>Kcnh4; |
|  |  |  |  |  |  |  |  |  | Slc7a5;<br>Atp2a1;<br>Pkd1l2 |
| ion<br>transmembrane<br>transporter activity<br>(GO:0015075) | 581 | 8 | 2.43 | 3.3 | 3.17E-03 | Up | Down | Down | Slc12a5;<br>Slc5a3;<br>Atp7b;<br>Slc24a4;<br>Kcnh4;<br>Slc7a5;<br>Atp2a1;<br>Pkd1l2 |
| metal ion<br>transmembrane<br>transporter activity<br>(GO:0046873) | 310 | 7 | 1.29 | 5.41 | 3.60E-04 | Up | Down | Down | Slc12a5;<br>Slc5a3;<br>Atp7b;<br>Slc24a4;<br>Kcnh4;<br>Atp2a1;<br>Pkd1l2 |
| monovalent<br>inorganic cation<br>transmembrane<br>transporter activity<br>(GO:0015077) | 261 | 5 | 1.09 | 4.59 | 5.10E-03 | Up | Down | Down | Slc12a5;<br>Slc5a3;<br>Kcnh4;<br>Atp2a1 |
| postsynaptic<br>neurotransmitter<br>receptor activity<br>(GO:0098960) | 49 | 2 | 0.2 | 9.77 | 1.91E-02 | Up | Down | Down | Lypd1;<br>Chrm2 |
| transmembrane<br>transporter activity<br>(GO:0022857) | 701 | 8 | 2.93 | 2.73 | 9.41E-03 | Up | Down | Down | Slc12a5;<br>Slc5a3;<br>Atp7b;<br>Slc24a4;<br>Kcnh4;<br>Slc7a5;<br>Atp2a1;<br>Pkd1l2 |
| ammonium ion<br>binding<br>(GO:0070405) | 80 | 2 | 0.33 | 5.99 | 4.59E-02 | Up | Down |  | Lypd1;<br>Chrm2 |
| calcium ion<br>transmembrane<br>transporter activity<br>(GO:0015085) | 84 | 3 | 0.35 | 8.55 | 5.74E-03 | Up | Down |  | Slc24a4;<br>Atp2a1;<br>Pkd1l2 |
| potassium ion<br>transmembrane<br>transporter activity<br>(GO:0015079) | 139 | 3 | 0.58 | 5.17 | 2.15E-02 | Up |  | Down | Slc12a5;<br>Kcnh4 |
| transporter activity<br>(GO:0005215) | 782 | 9 | 3.27 | 2.76 | 5.65E-03 | Up |  | Down | Slc12a5;<br>Kcnh4;<br>Pkd1l2 |
| extracellular<br>matrix structural<br>constituent<br>(GO:0005201) | 11 | 1 | 0.05 | 21.76 | 4.88E-02 | Up |  | Up | Fbn1 |
| oxidoreductase<br>activity, acting on<br>single donors with<br>incorporation of<br>molecular oxygen,<br>incorporation of<br>two atoms of<br>oxygen<br>(GO:0016702) | 10 | 1 | 0.04 | 23.94 | 4.48E-02 | Up |  | Up | Ido1 |
| DNA-binding<br>transcription factor | 618 | 2 | 0.31 | 6.55 | 3.60E-02 |  | Down | Down | Hif3a;<br>Egr3 |

A further analysis examining the molecular function of pathways that were changed across all groups is shown in Table 2. Here, pathways that were identified as significantly up- or down-regulated in comparisons between control and each of the three treatment groups are shown. For example, G protein-coupled receptor activity (GO:0004930) was down-regulated by DEX but showed a rebound increase. Cation transmembrane transporter activity (GO:0008324) on the other hand was upregulated by DEX and then reduced during the recovery period.

We assessed differentially expressed gene sets for regulator regions using I-cis (Imrichová *et al.* 2015). Consistent with induction by GC, we found enrichment for genes associated with binding sites for *Nr3c1* (the glucocorticoid receptor) (normalised enrichment score 5.48, >3 deemed significant). A stronger association was identified between H3K27me3, a marker of heterochromatin (NES 13.8), and those genes up-regulated in our model by chronic DEX treatment. Interestingly, we did not find enrichment for GR in the genes suppressed by DEX, this might be as the algorithm is less effective at identifying the distal enhancers associated with DEX suppression (McDowell *et al.* 2018). Instead, we found enrichment for CHD2 (a chromatin helicase), and H3K36me3 (which usually marks gene bodies, but can also be found in heterochromatin) (NES 12.1 and 11.8, respectively).

## METHODS

Studies were performed according to the Animals (Scientific Procedures) Act 1986 following specific approval from the UK Home Office (Project Licence P09E1F821), following review by the University of Edinburgh Animal Research Ethics Committee and in compliance with EU directive 2010/63/EU. Data are presented from three experiments (Figure 1). Experiments 1 and 2 used male 8-week-old C57Bl6/J mice (Jackson Laboratories, UK). Experiment 3 used *Pomc*-eGFP (Pinto *et al.* 2004) transgenic mice maintained on a C57Bl6/J background and bred in-house. Mice were acclimatised to controlled lighting conditions in the animal facility for two weeks prior to each experiment, at constant temperature (22°C). Animals were fed standard chow supplemented with sunflower seeds.

### Experiment 1

Mice (C57Bl6/J, n = 36) were housed 6 per cage with lights on 07:00-19:00, and then randomly assigned to receive dexamethasone (DEX) (~0.4 mg/kg/day)(Sigma Aldrich, UK) in drinking water, or vehicle for 4 weeks (Figure 1A). This dose of DEX would be the allosteric equivalent of 2.5 mg for a 70 kg human, the equivalent to around 16 mg of prednisolone. This ‘moderate’ dose sits in the middle of the range of a weaning dose of 40 mg down to 5 mg over 8-12 weeks commonly used to treat flares of rheumatoid arthritis or inflammatory bowel disease. Animals were weighed three times per week. Following 4 weeks treatment, DEX was withdrawn and animals were maintained in standard conditions thereafter. At withdrawal of DEX (time = 0), one week and four weeks, six animals per group, selected randomly, 2 per cage were sacrificed by cervical dislocation between 9 and 11am (early rest period). Whole pituitary, hypothalamus, and adrenal glands were dissected and frozen on dry ice and then stored at −80°C. Adrenal glands were weighed. Trunk blood was obtained and serum separated by centrifugation at 2000 *x g* for 10 minutes at 4°C and stored at −20°C. Glucose was measured using an Accuchek Aviva glucometer (Roche, UK).

### Experiment 2

Mice (C57Bl6/J, n = 48) were randomly assigned to four groups and were housed 3 animals per cage: Group A (control) received drinking water for 8 weeks; Group B (0 weeks recovery) received standard drinking water for 4 weeks and then DEX for 4 weeks; Group C (1 week recovery) received standard drinking water for 3 weeks, then DEX for 4 weeks with a final week of recovery; Group D (4 weeks recovery) receiving DEX in drinking water as above for four weeks with a switch to standard water for the following four weeks (Figure 1B). Two weeks prior to the start of the experiment lighting was reversed (lights on 21:00, lights off 09:00), otherwise conditions were as experiment 1. At the end of the experiment mice were randomly assigned to receive 30 minutes of restraint stress 20-60 minutes prior to sacrifice. As there was no significant effect of stress on measurements taken, these two groups have been combined. Mice were sacrificed by cervical dislocation and trunk blood collected into cold tubes prepared with 5 μL of 5% EDTA and placed on ice. Plasma was obtained by centrifugation as above and stored at −80°C until analysis. Anterior pituitary, hypothalamus, and adrenal were dissected and frozen on dry ice and stored at −80°C. All animals were sacrificed between 9:40 and 11:45 am (early wake period).

### Experiment 3

*Pomc*-eGFP mice (n = 36) were randomly assigned to the four groups described in experiment 2, three animals per cage with lights on 07:00-19:00 (Figure 1C). At the end of the experimental protocol, mice were sacrificed by cervical dislocation between 9:35 and 10:30am (early rest period). Plasma, hypothalamus, adrenal, and pituitaries were harvested as above. The posterior and intermediate pituitary lobes were carefully dissected and anterior pituitaries were dissociated as previously (Duncan *et al.* 2015). The early time point was chosen here to fit with other models being established in our laboratory. Briefly, pooled tissues, each pool from three mice, were minced with a razor blade before being placed into 2.5 ml DMEM with 25 mM Hepes (LifeTechnologies) supplemented with 10 μl/ml DNAse I (Sigma) and 0.25% trypsin (Worthington). The pituitaries were digested for 20 minutes at 37°C with gentle shaking every 5 minutes. Digestion was terminated by allowing cells to settle, removing the supernatant and trituration of the cells in 1 ml DMEM supplemented with 50 μl soybean trypsin inhibitor (Sigma), 50 μl Aprotinin (100 Kallikrein units) (Sigma), and 5 μl DNAse I. Cells were passed through a 70 μm cell strainer that was washed with further inhibition solution. Cells were then centrifuged 10 minutes at 100g at room temperature, the supernatant was removed and the cells resuspended in DMEM supplemented with 4.5 g/l glucose with L-glutamine and 25 mM HEPES (Life technologies), 0.3% BSA, 1x ITS liquid media supplement (Sigma), 4.2 μg/ml fibronectin (Sigma), and 1x antibiotic/antimycotic solution (Sigma) (penicillin, streptomycin, and amphotericin B) and triturated gently before being transported to the FACS facility.

### Isolation of eGFP positive cells

Cells were resuspended in 500 μl PBS supplemented with 25 mM HEPES and 5 mM EDTA (FACS buffer) and passed through a 35 μm cell strainer that was washed with a further 200 μl of the FACS buffer. Draq7 was added as a vitality marker and cells sorted using a Sony SH800. Gates were established using WT pituitaries to avoid capturing eGFP negative ve cells and to select single cells. Sorted single cells were resuspended into low bind Eppendorf tubes into 300 μl Trizol and were frozen on dry ice and stored at −80°C until sending for sequencing.

### RNA Extraction, QC, Library Preparation, and Illumina Sequencing

RNA isolation, library preparation, and sequencing reactions were conducted at GENEWIZ, LLC. (South Plainfield, NJ, USA). Total RNA was extracted from FACS sorted cells using Qiagen RNeasy Plus Universal Mini kit followed by manufacturer’s instructions (Qiagen, Hilden, Germany). The RNA samples were quantified using Qubit 2.0 Fluorometer (Life Technologies, Carlsbad, CA, USA) and were below the limit of detection.

Ultra-low input RNA sequencing library preparation used the Clontech SMART-V4 kit for cDNA Synthesis from 10 pg-10 ng total RNA and polyA amplification. Illumina Nextera XT kit was used to prepare the final DNA libraries according to manufacturer’s instructions. Integrity of the sequencing library was assessed on the Agilent TapeStation (Agilent Technologies, Palo Alto, CA, USA), and quantified by using Qubit 2.0 Fluorometer (Invitrogen, Carlsbad, CA) as well as by quantitative PCR (KAPA Biosystems, Wilmington, MA, USA).

The sequencing libraries were clustered on one lane of a patterned flowcell. After clustering, the flowcell was loaded on the Illumina HiSeq 4000 or equivalent instrument in High Output Mode according to manufacturer’s instructions. The samples were sequenced using a 2×150 Paired End (PE) configuration. Image analysis and base calling were conducted by the HiSeq Control Software (HCS). Raw sequence data (.bcl files) generated from Illumina HiSeq were converted into fastq files and de-multiplexed using Illumina's bcl2fastq 2.17 software. One mismatch was allowed for index sequence identification.

### Steroid quantification

For experiment 1, 11-dehydrocorticosterone (A) and corticosterone (B) were measured by tandem liquid chromatography mass spectrometry (LC-MS/MS) in 30 μL of serum at the mass spectrometry core facility, University of Edinburgh. Briefly, plasma (30 μL) was enriched with internal standards (D4-cortisol (D4F) and epi-corticosterone ((Epi-B), 2.5 ng). Chloroform (10:1) was added and vortexed. The supernatant was reduced to dryness under oxygen-free nitrogen (OFN) at 60°C and reconstituted in water:acetonitrile(70 μL; 70:30 (v/v)). Samples were extracted alongside a calibration curve of A and B with D4F and epi-B as internal standards. Quantitative analysis of the extracts was carried out on a Shimadzu Nexera – Sciex QTrap 6500+ LC-MS/MS instrument, operated in positive ion mode, as described previously (Nixon *et al.* 2016; Verma *et al.* 2019).

For experiment 2, a low volume automated extraction method was developed for mouse plasma and combined with an LC-MS/MS method adapted from (Agnew *et al.* 2019), to detect corticosterone, 11-dehydrocorticosterone, and dexamethasone by LC-MS/MS in only 10 μL of mouse plasma. Briefly, 10 μL of plasma was diluted to 100 μL with water, prepared alongside a calibration curve of the steroids of interest. The plasma, calibration standards, and blanks were dispensed manually as aliquots (100 μL) into individual wells of a 2 mL 96 deep-well polypropylene plate (Waters, UK). An internal standard solution of isotopically labelled standards (d8-corticosterone (Cambridge isotope laboratories), and d4-dexamethasone (Sigma-Cerrilliant)) in methanol was added (20 μL; 10 ng) to each well except for the double blanks. The plate was agitated on a plate shaker (2 minutes) and transferred to a Biotage® Extrahera™ automated sample processor (Biotage, Uppsala, Sweden) where formic acid (100 μL, 0.1%, v/v) was added to each well. Samples were incubated at room temperature (18-22°C; 5 minutes) and transferred to an SLE+ 200 plate by the robot, loaded onto the SLE material under positive pressure using compressed air. Following a wait (5 minutes) the analytes were eluted from the SLE material into a deep-well collection plate by positive pressure following the addition of dichloromethane/propan-2-ol (98:2; 4x 450 μL). The eluate was dried down under a stream of heated oxygen free nitrogen (OFN, 40°C) on an SPE DryTM Dual Sample Concentrator System (Biotage, Uppsala, Sweden). Once dry the extracts were dissolved in water/methanol (70:30, 100 μL), the plate was sealed then shaken on a plate shaker (10 minutes) and sealed with a zone-free plate seal, before injecting directly from the 96-well plate for LC-MS/MS analysis.

Samples were injected (10 μL) onto a Kinetex C18 column (150 × 3 mm; 2 μm, Phenomenex, Macclesfield, UK) at 40°C using a flow rate of 0.5 mL/min, mobile phase A – water 0.05 mM ammonium fluoride, B – methanol 0.05 mM ammonium fluoride from 55 to 100% B over 16 minutes, protected by a Kinetex KrudKatcher (Phenomenex, Macclesfield, UK), followed by analysis on a QTrap 6500+ Linear Ion Trap Quadrupole Mass Spectrometer (AB Sciex, UK) system TurboIonspray source operated at 600°C. A, B, and dexamethasone were detected with the following transitions; *m/z* 345.1.1 → 121.2, *m/z* 347.1 → 121.1, *m/z* 393.1 → 373.2 while internal standards d8-corticosterone and d4-dexamethasone were detected with the following transitions; *m/z* 355.3 → 125.1 and *m/z* 397.1 → 377.2.

The peak areas of each steroid and each internal standard were integrated using Quantitate in Analyst 1.6.3. Linear regression analysis of calibration standards, calculated using peak area ratios of steroid of interest to internal standard, was used to determine the concentration of the steroid of interest in the samples. *R* _2_ > 0.99 was considered acceptable and within each batch of samples the accuracy at the upper and lower limits were only accepted if accuracy <20%. The amount of steroid was calculated using linear regression analysis of the peak area ratio.

### ACTH quantification

ACTH was measured in 25 - 50 μl of plasma using an ACTH ELISA kit M046006 (MD biosciences, Switzerland) according to manufacturers’ instructions. The assay has low cross-reactivity with other products of *Pomc*. Samples were diluted 1:4 or 1:8. The assay had a lower limit of detection of 4 pg/ml, has an intra-assay precision (CV) of 6.7% and an inter-assay precision (CV) of 7.1%.

### qRT-PCR

RNA was extracted from tissue using the Reliaprep miniprep system (Promega, UK) according to the standard protocol except that treatment with DNase to remove genomic DNA was extended for 30 minutes. RNA purity was assessed using spectrophotometry to ensure 260/280 readings were >1.8. RNA integrity for subsets of samples was assessed by inspection of RNA on a 1% agarose gel. RNA was reverse transcribed using Superscript IV (Thermo-Fisher, UK) in accordance with manufacturer’s instructions with random hexamers and oligodT (50:50). qRT-PCR was conducted using an ABI 7500 and relative expression determined, with stably expressed reference genes determined using NormFinder: for experiment 1, adrenal reference genes were *Gapdh* and *Ipo8*; for pituitary, *Ppia* and *Ipo8*; and for hypothalamus, *Ppia* and *Gapdh*. For experiment 2 (adrenal) *Gapdh* and *Fbox10* were used as house keeping genes. Data are presented as delta-delta Ct versus the control; in experiment 1 the control from the same time point and in experiment 2 with the control arm (group A).

### Statistics

Differences between groups were assessed by linear mixed model. Normality was assessed by visualisation of q-q plots. For experiment 1, treatment and time were used as dependent variables and cage as a random variable. For experiments 2 and 3, group was used as the dependent variable and cage as a random factor. For assessing bodyweight, group and time were used as dependent variables and individuals within cage as random factors to allow for repeated measures. For ease of comparison of qPCR experiments and quantification of glucocorticoids, data from experiment 1 were further analysed grouping together results from the different times in the control arm (delta Ct still relative to the time point control). Where a significant interaction between time and treatment was identified (experiment 1), or an effect of group (experiment 1, 2, and 3), pairwise comparison was made, corrected for using Tukey’s HSD. Statistical analysis was conducted using R version 3.6.2; p<0.05 was taken as statistically significant. Data are referred presented in box and whisker plots (with median and interquartile range).

### RNA-seq bioinformatic analysis

Nextera adapters were removed from paired end FASTA files using trimgalore v0.6.5 using the flags –paired and –2colour. Quality control, including successful removal of adapters, assessment of phred scores and sequence length were assessed using FastQC v0.11.9. Trimmed reads were aligned to the mouse genome (GRCm38 release 98) using STAR v2.7.1a. Aligned reads were quantified using featureCounts v2.0.0 using the flags -p (paired end), -Q 20 (mapping quality >20) and -s 2 (reverse stranded) (Liao *et al.* 2014). Count files were then analysed by EdgeR keeping genes with >0.1 counts per million (cpm) in at least three samples and with differential expression based upon trended dispersion. For comparison with qRT-PCR data (Fig. 3C), cpm relative to the control group (A) were calculated for each gene. 17.5 million to 27.1 million reads were obtained from each sample of corticotrophs pooled from three pituitaries (Fig. S2C).

## DISCUSSION

We exposed mice to the synthetic GR agonist, DEX for four weeks in drinking water and show sustained suppression of the HPA axis persisting for at least one week following treatment withdrawal. We showed that at one week following DEX withdrawal, some animals show a compensatory increase in plasma ACTH and corticotroph *Pomc* expression whilst others still demonstrate suppressed corticotroph action given the relative corticosterone deficiency. DEX has a significant effect on the corticotroph transcriptome; whilst one week following treatment withdrawal, the majority of genes affected by DEX have returned to control levels, a number of genes do retain differential expression that is persistent at least 1 week after withdrawal of DEX, and which may play a role in disrupted activity.

As expected, we found evidence for reduced adrenal function persisting one week following withdrawal of DEX treatment (Finco *et al.* 2018; Spiga *et al.* 2020). A non-significant but persistent reduction in adrenal size and a significant reduction in the abundance of transcripts for genes regulating steroidogenesis was found one week following DEX withdrawal. Glucocorticoid-producing cells of the zona fasciculata are generated from *Gli1* and *Shh* expressing progenitor cells both in response to stress (Steenblock *et al.* 2018), and recovery from withdrawal of exogenous glucocorticoid treatment (Finco *et al.* 2018). The signals driving adrenal recovery in this paradigm are unclear; whilst ACTH certainly drives hypertrophy of the zona fasciculata (Baccaro *et al.* 2007), other endocrine and paracrine factors (e.g. sympathetic stimulation (Steenblock *et al.* 2018) and VEGF signalling (Mallet *et al.* 2003)) also play a role in driving recovery. In mice, co-treatment with ACTH attenuates DEX-induced adrenal atrophy further, highlighting the trophic role of this hormone in the adrenal (Mallet *et al.* 2003).

The adrenal response to ACTH is reduced both in humans and in rodent models following withdrawal of chronic exogenous GC treatment (Borresen *et al.* 2017; Spiga *et al.* 2020). Previous studies highlight normalised ACTH plasma measurements in the context of ongoing adrenal insufficiency, suggesting an ongoing defect at the level of the adrenal (Schuetz *et al.* 2008; Borresen *et al.* 2017; Spiga *et al.* 2020). However, another interpretation of these data is that ACTH levels should be increased in the milieu of relative adrenal insufficiency and thus normalised ACTH levels are suggestive of ongoing attenuated corticotroph (or hypothalamic) function. Further, that expression of the steroidogenic enzymes that remain suppressed one week following DEX withdrawal are driven by ACTH (Ruggiero & Lalli 2016) might also point towards ongoing impaired activity at the higher HPA axis. Taken together, data suggest that exogenous GC exposure suppresses both adrenal and pituitary function, and confirm that recovery of adrenal function is dependent (at least in part) upon recovery of the pituitary.

One week following withdrawal of DEX, we find heterogeneity in the HPA axis activity of mice. In the waking phase (pm) we show that whilst 5 of 9 mice exhibit the expected increase in ACTH associated with impaired adrenal function (consistent with recovery of corticotroph activity), 2 of 9 have ACTH that is inappropriately low given the low corticosterone levels. We see a similar pattern in the transcriptome of isolated corticotrophs where 1 pool out of 3 shows low (similar to control) levels of *Pomc* transcript when this should be elevated in the context of adrenal insufficiency. This variability is reminiscent of human studies where only a proportion of patients are found to have ongoing adrenal insufficiency following withdrawal of exogenous GC (e.g. 10% of patients at 6 months) (Dinsen *et al.* 2013; Joseph *et al.* 2016). The data suggest inter-individual differences in the trajectory of recovery of the mice in terms of ACTH release that could be established in further studies looking at further ‘snap shots’ of the recovery process following DEX withdrawal.

A number of possibilities could account for these differences: (1) we show variance in the levels of plasma DEX attained as a result of the exposure in drinking water, thus the variability could be attributed to different dose exposures as a result of differences in water intake as mice were co-housed; (2) social hierarchy may affect HPA axis activity as observed in Cynmolgus (Jimenez *et al.* 2017) (although we pooled from a cage each for each experiment); (3) finally, factors that influence GC sensitivity of the HPA axis, for example epigenetic changes in the hypothalamus as a result of early life or prenatal stress exposure (Ostrander *et al.* 2006; van Bodegom *et al.* 2017; Matthews & McGowan 2019), could account for inter-individual differences.

DEX at 0.4 mg/kg likely reaches lower concentrations within the blood brain barrier than plasma; 0.2 mg/kg/mouse reached brain concentrations of 30% that of plasma as a result of extrusion by Mdr1/Abcb1 (Schinkel *et al.* 1995), a p-glycoprotein that helps to reduce accumulation of toxins within the brain, and which exports DEX. Thus a relative GC deficient state was likely induced by our DEX exposure. We did not find changes in whole hypothalamus transcript levels for *Crh* or *Avp* when measured in the rest period. The magnocellular component (which does not contribute directly to HPA axis activity) in which these transcripts are also found abundantly may account for the lack of change here. As we have dissected whole hypothalamus we are unable to assess specific changes in the parvocellular neurones that regulate anterior pituitary hormone release.

We explored recovery at the level of the pituitary further by isolating corticotrophs from *Pomc*-eGFP mice. The eGFP profile of the isolated cells exhibited a bimodal distribution, most apparent in groups A and C; in groups B and D there was a shift in the population to the left with a greater proportion of ‘low eGFP’ expressing cells.

Two possibilities to account for multiple populations of cells arise here. First, inclusion of melanotrophs that also produce large quantities of *Pomc*. These cells were removed by careful dissection of the *pars intermedia*, but some melanotrophs may be present in the anterior pituitary (Mayran *et al.* 2019). We did not find any association between expression of *Pcsk2* and *Pax7*, markers of melanotrophs, and those samples with larger populations of eGFP ‘high’ cells. Further our own experience with dispersed GFP +ve cells *in vitro* suggests that <1% of cells fail to respond to CRH/AVP, a mechanism specific to corticotrophs. Given that in some pools of cells, >30% of cells made up those in the eGFP ‘high’ peak, it is hard to reconcile that these cells all made up melanotrophs of the anterior pituitary. However, we cannot discount the fact that the population may have included some melanotrophs. Second, that corticotroph cells can exist in two ‘states’. To date single cell RNA-seq from pituitaries suggests a spectrum of transcriptomic activity (as assessed by levels of *Pomc* transcript) in cells identified as corticotrophs (Cheung *et al.* 2018). Variable transcription rates across a population are reminiscent of those described from live cell imaging of pituitary lactotrophs (Featherstone *et al.* 2016). Thus a heterogenous population of corticotroph cells may have been shifted ‘left’ to form a more homogenous population by treatment with DEX, leaving the possibility that the pituitary has undergone physiological change that has not been detected by basal ACTH and corticosterone measurement.

As previous *in vitro* studies had shown a persistent stimulation of *Fkbp5* mRNA expression following withdrawal of DEX (Jubb *et al.* 2017), we wished to assess if the chronic DEX treatment in mice resulted in sustained transcriptional changes in the corticotrophs that might explain the delay in recovery of the HPA axis. Only 1 gene was supressed by DEX and exhibited ongoing suppression either 1 or 4 weeks following treatment withdrawal. 2 of the 101 genes up-regulated by DEX remained elevated 4 weeks after withdrawal of the DEX, with 5 genes showing ongoing upregulation 1 week after DEX withdrawal. We thus found little evidence to support persistent global dysregulation of gene expression as a result of DEX exposure, but we did find a small number of genes with persistent differential expression, at least at 1 week after stopping the DEX

We identified several genes with differential expression 1 week after withdrawal of DEX that may play a role in regulation of corticotroph activity. Two G-protein-coupled receptors with known significance in corticotroph biology were reduced by DEX exposure, but showed significant increase during the recovery process. *Sstr2*, the somatostatin 2 receptor is a target of Pasireotide, which is being trialled for use in treatment of Cushing’s disease. The change in expression of *Sstr2* as a result of GC exposure could have implications for the efficacy of this treatment, and selection of patients most likely to benefit (Lacroix *et al.* 2020). *Aplr*, the apelin receptor is of interest as apelin is a stimulator of ACTH release and steroidogenesis (Newson *et al.* 2009; Yang *et al.* 2019). This effect was previously attributed to a paracrine effect within the hypothalamus but these data suggest a direct role on corticotroph function is possible, which may affect signalling even one month following withdrawal of therapy. The transporter *Slc12a5*, (also known as KCC2) was also up-regulated by DEX, the channel is inhibited by loop diuretics, and *in* vitro has been shown to inhibit ACTH release (Heisler 1991). Further transporter/ion channel proteins were identified with hitherto unknown function in corticotrophs. These may make novel targets for limiting ACTH release in patients with Cushing’s, but also potential pathways that could be activated to try to increase the rate of recovery in patients with GC-induced adrenal insufficiency or disordered HPA axis regulation. We also identified a collection of kallikrein genes up-regulated by DEX. Previous experiments have shown small reductions in ACTH release in response to hypoglycaemia where rats were co-treated with a kallikrein inhibitor (Madeddu *et al.* 1992); thus, this is a potential mechanism further inhibiting ACTH release in response to GC. A number of genes associated with extra-cellular matrix were identified, *Fbn1, Col3a1*, and *Mfap4*. Given persistent changes seen in some of these genes following DEX withdrawal, it is interesting to postulate that the intercellular network of corticotophs may have been affected by DEX, as has been seen in lactotrophs following 1st lactation (Hodson *et al.* 2012).

There are a number of limitations to our study. The number of timepoints we have studied has been limited by resources, and thus we have only looked at a small number of ‘snap shots’ during the recovery process. The dynamics of recovery between these time points, and if these exhibit inter-individual differences would be important foci for future work. Further, as we required to pool pituitaries in order to obtain sufficient numbers of corticotrophs for RNA-sequencing, our data lack resolution at the individual level. Mice likely experienced stress at time of sacrifice, and whilst procedures were kept consistent for all experiments, perceived stress could account for some of the variation in hormone (especially ACTH) measurements.

To conclude, we established a model of chronic glucocorticoid treatment in mice and found that 28 days of DEX exposure had suppressed HPA activity, which persisted 7 days following treatment withdrawal. DEX treatment had a persistent effect on adrenal steroidogenesis and abundance of adrenal transcripts for steroidogenic enzymes that lasted at least 7 days. DEX suppressed corticotroph *Pomc* transcription, an effect which had recovered in some animals one week following withdrawal but persisted in others, mirroring the pattern seen in ACTH measurement. Earlier time points during recovery and an increase the number of individuals would be useful for future experiments examining regulation of suppression, and recovery of hypothalamic and corticotroph activity following chronic GC exposure. A persistent but small change in the transcriptome of *Pomc-*expressing isolated GFP positive cells was identified 4 weeks following DEX withdrawal. This was not associated with altered corticosterone or ACTH production, but might affect further pituitary response to other stimuli, for example repeat steroid prescription or stress, and could contribute to the longevity of HPA axis suppression seen in some humans following withdrawal of chronic GC treatment.

## Acknowledgements

We are very grateful to insightful discussion with Karen Chapman, Jacques Drouin, and Sean Bankier. We acknowledge the financial support of NHS Research Scotland (NRS), through Edinburgh Clinical Research Facility for the Mass Spectrometry Core. Studies were supported by MRC project grant to MJS and Wellcome ISSF3 grant to TJGC. BRW is a Wellcome Investigator.

## Figure legends

**Figure S1.**
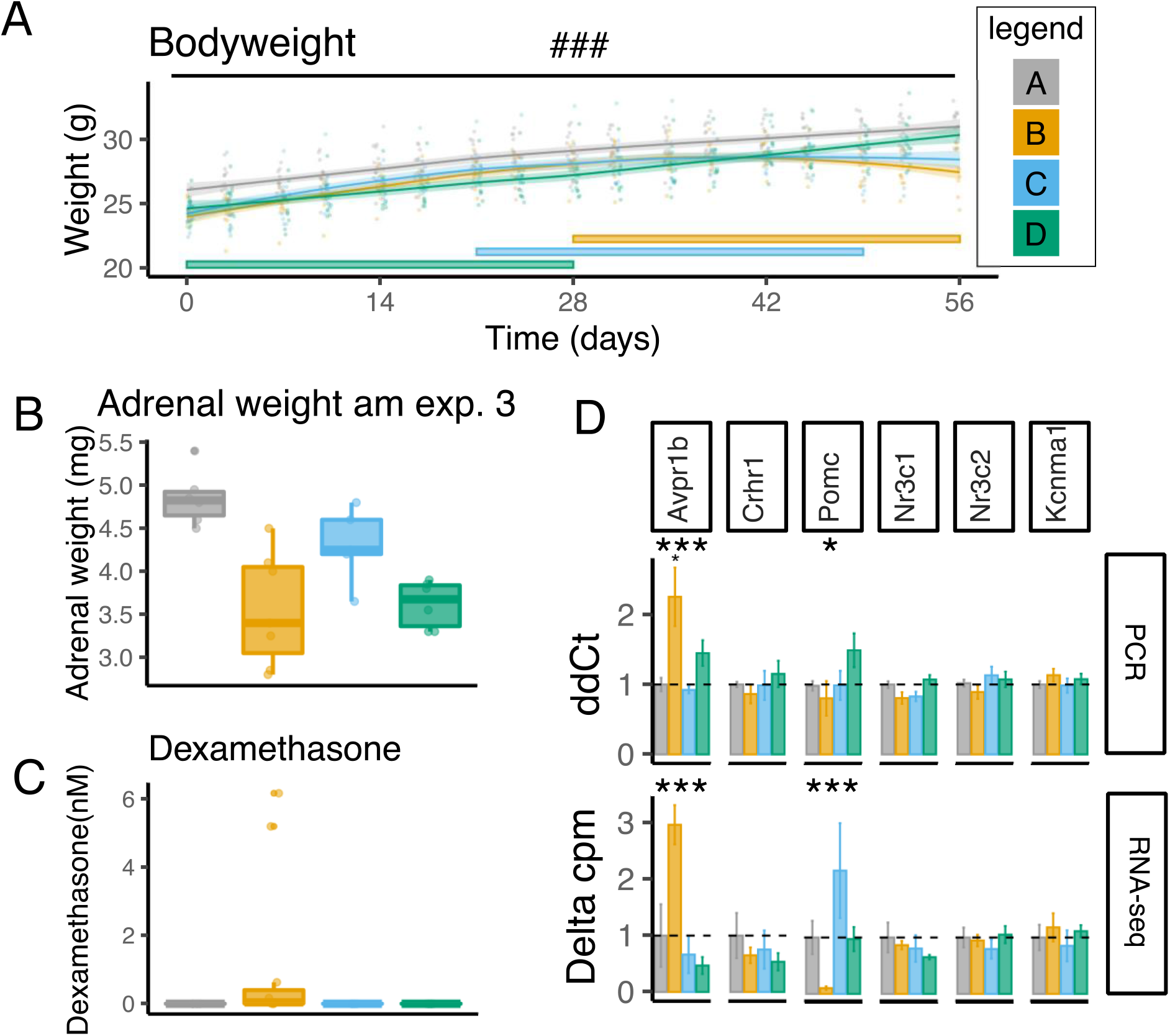
**A. A. Bodyweight**. DEX treat inhibited weight gain but this returned to control levels 4 weeks after stopping treatment. The weight of each mouse in experiment 2 is shown as representative. Mice were weighed twice a week. Mean and 95% confidence intervals are presented as lines. N = 16 cages of 3 mice. ### p<0.0001 for interaction between weight and time assessed by linear mixed model with cage and individual as random factor to account for repeated measures. Coloured bars at the bottom of the figure indicate time when mice were exposed to DEX. **B. Adrenal weight** weights of adrenals from experiment 3 were not significantly affected by treatment but were reduced and remained lower 4 weeks after stopping DEX. **C. Dexamethasone levels**. Plasma dexamethasone was measured at the end of the rest period (experiment 2). DEX increased measured DEX, which were undetectable in control groups or following 1 week of withdrawal of DEX. **D. Comparison of RNA-seq and qPCR from whole pituitary.** qPCR of whole pituitary and 6 key genes for corticotroph function is shown in the top panel. Pituitaries were collected in experiment 1 (n = 5-6 from 3 cages). Data analysed by linear mixed model with group as dependent variable and cage as random factor. Tukey-adjusted post hoc tests compared to control group are indicated above bars (small asterixis) where significant differences were identified. *** p<0.001, ** p<0.01, * p<0.05. The bottom panel shows for comparison the relative cpm to Group A (control) from the RNA-seq (experiment 3) (n = 3 pools of 2-3 pituitaries). *** p<0.0001; FDR. **Legend.** Grey dots and boxes represent control mice (Group A), yellow those who have had 4 weeks of DEX treatment (Group B), blue those one week after withdrawal of DEX (Group C), and green those 4 weeks after treatment withdrawal (Group D). Data analysed by linear mixed model with group as dependent variable and cage as random factor.

**Figure S2.**
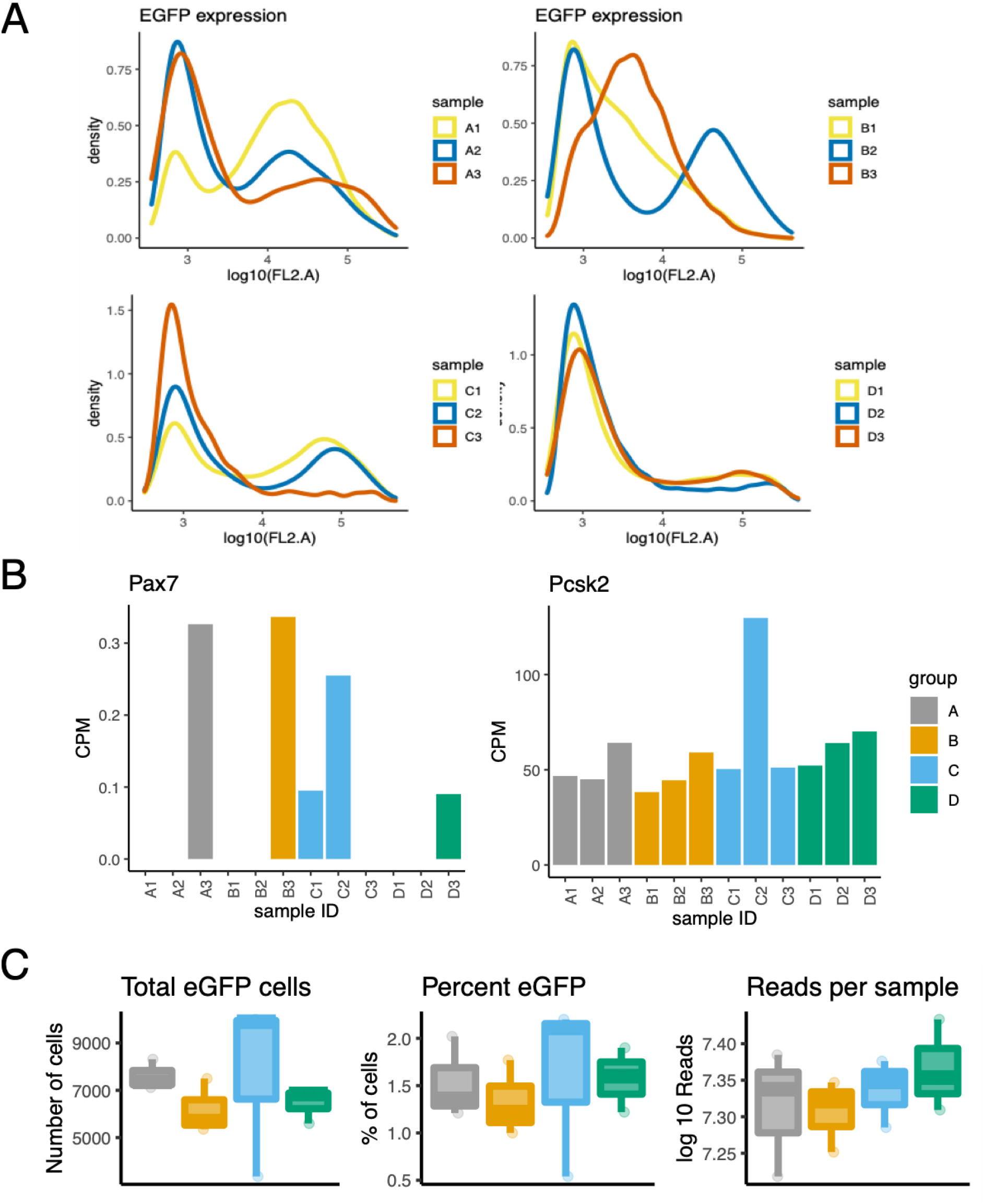
**A. Fluorescence of isolated cells from each sort.** FACS GFP distribution is shown for each sort of 2-3 anterior pituitaries following treatment (experiment 3). **B.** Raw counts of Pax7 and Pcsk2, markers of melanotrophs for comparison with panel A. Expression of melanotroph specific genes does not associate with sorts with larger secondary peaks of higher fluorescence. **C.** Number of cells, percent of fluorescent cells, and number of reads obtained form each sample from dissociated anterior pituitaries obtained in experiment 3. There were no significant difference between groups. N = 3 pools of 2-3 pituitaries. **Legend.** Grey dots and boxes represent control mice (Group A), yellow those who have had 4 weeks of DEX treatment (Group B), blue those one week after withdrawal of DEX (Group C), and green those 4 weeks after treatment withdrawal (Group D).

**SUPPLEMENTAL TABLE 1:** Gene ontology analysis of GC induced genes (Molecular function)

| Enrichment FDR | Genes in list | Total genes | Functional Category | Genes |
| --- | --- | --- | --- | --- |
| 0.000 | 7 | 135 | Hydrolase activity, acting on carbon-nitrogen (but not peptide) bonds | MTHFD2 KLK1B16 KLK1B26 KLK1B24 KLK1B5 KLK1B4 KLK1B21 |
| 0.000 | 6 | 83 | Hydrolase activity, acting on carbon-nitrogen (but not peptide) bonds, in linear amides | KLK1B16 KLK1B26 KLK1B24 KLK1B5 KLK1B4 KLK1B21 |
| 0.001 | 7 | 198 | Serine-type endopeptidase activity | KLK1B5 PRSS57 KLK1B16 KLK1B26 KLK1B24 KLK1B4 KLK1B21 |
| 0.002 | 7 | 217 | Serine-type peptidase activity | KLK1B5 PRSS57 KLK1B16 KLK1B26 KLK1B24 KLK1B4 KLK1B21 |
| 0.002 | 7 | 222 | Serine hydrolase activity | KLK1B5 PRSS57 KLK1B16 KLK1B26 KLK1B24 KLK1B4 KLK1B21 |
| 0.003 | 3 | 23 | Structural constituent of muscle | MYOM1 MYH11 KRT19 |
| 0.004 | 9 | 469 | Endopeptidase activity | ATG4A KLK1B5 PRSS57 KLK1B16 KLK1B26 ADAMTS14 KLK1B24 KLK1B4 KLK1B21 |
| 0.004 | 11 | 683 | Peptidase activity | ATG4A KLK1B26 ADAMTS14 KLK1B5 KLK1B21 PRSS57 USP54 KLK1B16 KLK1B24 KLK1B4 ADAMTSL3 |
| 0.006 | 2 | 7 | Netrin receptor activity | UNC5B UNC5A |
| 0.009 | 8 | 427 | Metal ion transmembrane transporter activity | ATP7B SLC12A5 ATP2A1 KCNH4 SLC24A4 SLC5A3 ASIC2 TMEM37 |
| 0.009 | 10 | 654 | Peptidase activity, acting on L-amino acid peptides | ATG4A KLK1B5 PRSS57 USP54 KLK1B16 KLK1B26 ADAMTS14 KLK1B24 KLK1B4 KLK1B21 |
| 0.014 | 2 | 12 | Diacylglycerol kinase activity | DGKI DGKK |
| 0.020 | 9 | 622 | Cation transmembrane transporter activity | ATP7B SLC12A5 ATP2A1 KCNH4 SLC24A4 SLC5A3 SLC44A4 ASIC2 TMEM37 |
| 0.030 | 2 | 19 | NAD+ kinase activity | DGKI DGKK |
| 0.038 | 8 | 574 | Inorganic cation transmembrane transporter activity | ATP7B SLC12A5 ATP2A1 KCNH4 SLC24A4 SLC5A3 ASIC2 TMEM37 |
| 0.040 | 10 | 851 | Ion transmembrane transporter activity | ATP7B SLC12A5 ATP2A1 KCNH4 SLC7A5 SLC24A4 SLC5A3 SLC44A4 ASIC2 TMEM37 |
| 0.045 | 6 | 362 | Monovalent inorganic cation transmembrane transporter activity | SLC12A5 ATP2A1 KCNH4 SLC24A4 SLC5A3 ASIC2 |

**SUPPLEMENTAL TABLE 2:** Gene ontology analysis of GC suppressed genes (Molecular function)

| Enrichment FDR | Genes in list | Total genes | Functional Category | Genes |
| --- | --- | --- | --- | --- |
| 0.001 | 3 | 51 | Antigen binding | H2-Q6 H2-AA H2-AB1 |
| 0.001 | 5 | 291 | Peptide binding | SSTR2 H2-Q6 H2-AA H2-AB1 CD74 |
| 0.001 | 3 | 33 | Peptide antigen binding | H2-Q6 H2-AA H2-AB1 |
| 0.001 | 2 | 7 | CD4 receptor binding | IL16 CD74 |
| 0.001 | 5 | 358 | Amide binding | SSTR2 H2-Q6 H2-AA H2-AB1 CD74 |
| 0.001 | 2 | 12 | Platelet-derived growth factor binding | PDGFRB COL3A1 |
| 0.002 | 8 | 1443 | Protein dimerization activity | RRM2 PECAM1 TOP2A ITGAL ADGRL4 H2-AA H2-Q6 H2-AB1 |
| 0.005 | 8 | 1628 | Signaling receptor binding | POMC H2-Q6 IL16 CD74 PDGFRB COL3A1 KDR NMB |
| 0.005 | 3 | 144 | Growth factor binding | PDGFRB COL3A1 KDR |
| 0.005 | 5 | 607 | Protein heterodimerization activity | TOP2A ITGAL H2-AA H2-Q6 H2-AB1 |
| 0.005 | 2 | 34 | Neuropeptide receptor binding | POMC NMB |
| 0.013 | 3 | 223 | Cell adhesion molecule binding | COL3A1 ITGAL KDR |
| 0.014 | 2 | 68 | Transmembrane receptor protein tyrosine kinase activity | PDGFRB KDR |
| 0.014 | 6 | 1207 | Protein-containing complex binding | NCKAP1L CD74 COL3A1 ITGAL KDR H2-Q6 |
| 0.020 | 2 | 85 | Transmembrane receptor protein kinase activity | PDGFRB KDR |
| 0.023 | 8 | 2327 | Enzyme binding | NCKAP1L CCNB2 TOP2A PDGFRB COL3A1 CD74 H2-AB1 PECAM1 |
| 0.026 | 4 | 633 | Protein kinase activity | CCNB2 PBK PDGFRB KDR |
| 0.029 | 8 | 2473 | Signaling receptor activity | ADGRL4 APLNR SSTR2 CD74 PDGFRB ITGAL RTN4RL1 KDR |
| 0.030 | 2 | 118 | Integrin binding | COL3A1 KDR |
| 0.030 | 8 | 2521 | Molecular transducer activity | ADGRL4 APLNR SSTR2 CD74 PDGFRB ITGAL RTN4RL1 KDR |
| 0.031 | 2 | 128 | Extracellular matrix structural constituent | COL3A1 MFAP4 |
| 0.032 | 2 | 136 | Protein tyrosine kinase activity | PDGFRB KDR |
| 0.032 | 4 | 735 | Phosphotransferase activity, alcohol group as acceptor | CCNB2 PBK PDGFRB KDR |
| 0.032 | 2 | 140 | G protein-coupled peptide receptor activity | APLNR SSTR2 |
| 0.035 | 2 | 149 | Peptide receptor activity | APLNR SSTR2 |
| 0.039 | 2 | 162 | Protein kinase regulator activity | CCNB2 NCKAP1L |
| 0.043 | 4 | 840 | Kinase activity | CCNB2 PBK PDGFRB KDR |
| 0.044 | 7 | 2304 | Transmembrane signaling receptor activity | ADGRL4 APLNR SSTR2 CD74 PDGFRB ITGAL KDR |
| 0.050 | 2 | 194 | Kinase regulator activity | CCNB2 NCKAP1L |

